# zUMIs - A fast and flexible pipeline to process RNA sequencing data with UMIs

**DOI:** 10.1101/153940

**Authors:** Swati Parekh, Christoph Ziegenhain, Beate Vieth, Wolfgang Enard, Ines Hellmann

## Abstract

Single cell RNA-seq (scRNA-seq) experiments typically analyze hundreds or thousands of cells after amplification of the cDNA. The high throughput is made possible by the early introduction of sample-specific barcodes (BCs) and the amplification bias is alleviated by unique molecular identifiers (UMIs). Thus the ideal analysis pipeline for scRNA-seq data needs to efficiently tabulate reads according to both BC and UMI. *zUMIs* is such a pipeline, it can handle both known and random BCs and also efficiently collapses UMIs, either just for Exon mapping reads or for both Exon and Intron mapping reads. Another unique feature of *zUMIs* is the adaptive downsampling function, that facilitates dealing with hugely varying library sizes, but also allows to evaluate whether the library has been sequenced to saturation. *zUMIs* flexibility allows to accommodate data generated with any of the major scRNA-seq protocols that use BCs and UMIs. To illustrate the utility of *zUMIs*, we analysed a single-nucleus RNA-seq dataset and show that more than 35% of all reads map to Introns. We furthermore show that these intronic reads are informative about expression levels, significantly increasing the number of detected genes and improving the cluster resolution. **Availability:** https://github.com/sdparekh/zUMIs

## Introduction

The recent development of increasingly sensitive protocols allows to generate RNA-seq libraries of single cells [1]. The throughput of such single-cell RNA-sequencing (scRNA-seq) protocols is rapidly increasing, enabling the profiling of tens of thousands of cells [2, 3] and opening exciting possibilities to analyse cellular identities [4, 5]. As the required amplification from such low starting amounts introduces substantial amounts of noise [6], many scRNA-seq protocols incorporate unique molecular identifiers (UMIs) to label individual cDNA molecules with a random nucleotide sequence before amplification [7]. This enables the computational removal of amplification noise and thus increases the power to detect expression differences between cells [8, 9]. To increase the throughput, many protocols also incorporate sample-specific barcodes (BCs) to label all cDNA molecules of a single cell with a nucleotide sequence before library generation [10]. This allows for early pooling, which further decreases amplification noise [6]. Additionally, for cell types such as primary neurons it has been proven to be more feasible to isolate RNA from single nuclei rather than whole cells [11, 12]. This decreases mRNA amounts further, so that it has been suggested to count Intron mapping reads originating from nascent RNAs as part of single cell expression profiles [11]. However, the few bioinformatic tools that process RNA-seq data with UMIs and BCs have limitations (Table 1). For example the Drop-seq-tools is not open source [13]. While Cell Ranger is open, it is exceedingly difficult to adapt the code to new or unknown sample barcodes and other library types. Other tools are specifically designed to work with one mapping algorithm and focus mainly on transcriptomes [14, 15]. Furthermore, the only other UMI-RNA-seq pipeline providing the utility to also consider Intron mapping reads, dropEst [16], is only applicable to droplet-based protocols. Here, we present *zUMIs*, a fast and flexible pipeline that overcomes these limitations.

#### Key Points

- zUMIs processes UMI-based RNA-seq data from raw reads to count tables in one command.
- Unique features of zUMIs:
  – Automatic cell barcode selection
  – Adaptive downsampling
  – Counting of Intron mapping reads for gene expression quantification
- zUMIs is compatible with all major UMI-based RNA-seq library protocols.

**Table 1.** Features of available UMI pipelines for the quantification of gene expression data. We consider whether the pipeline is open source, has sequence quality filters for cell barcodes (BC) and UMIs, mappers, UMI-collapsing options, options for BC detection (A - automatically infer intact BCs, WL - extract only the given list of known BCs, top-n - order BCs according the number of reads and keep the top n BCs, EM - merge BCs with given edit-distance), whether it can count Intron mapping reads, whether it offers a utility to make varying library sizes more comparable via downsampling and finally with which RNA-seq library preparation protocols it is compatible.

| Name | Reference | Open Source | Quality filter | UMI collapsing | Mapper | BC detection | Intron | Down-sampling | Compatible UMI library protocols |
| --- | --- | --- | --- | --- | --- | --- | --- | --- | --- |
| Cell Ranger | [2] | yes | BC+UMI | Hamming distance | STAR | A | no | yes | [2] |
| CEL-seq | [15] | yes | BC+UMI | identity only | bowtie2 | WL | no | no | [44, 15] |
| dropEst | [16] | yes | BC | frequency-based | TopHat2 or Kallisto | WL,top-n,EM | yes | no | [13, 19, 2] |
| Drop-seq-tools | [13] | no | BC+UMI | Hamming distance | STAR | WL,top-n | no | no | [13, 17, 15] |
| scPipe | [45] | yes | BC+UMI | Hamming distance | subread | WL,top-n | no | no | [44, 18, 17, 13] |
| umis | [14] | yes | BC | frequency-based | Kallisto | WL,top-n,EM | no | no | [17, 44, 46, 18, 13, 19, 2] |
| UMI-tools | [25] | yes | BC+UMI | network-based | BWA | WL | no | no | [17, 19] |
| zUMIs | This work | yes | BC+UMI | Hamming distance | STAR | A,WL,top-n | yes | yes | [17, 44, 46, 18, 13, 15, 21, 12, 3, 2] |

## Findings

*zUMIs* is a pipeline to process RNA-seq data that were multiplexed using cell barcodes and also contain UMIs. Read pairs are filtered to remove reads with low quality BCs or UMIs based on sequence and then mapped to a reference genome (Figure 1). Next, *zUMIs* generates UMI and read count tables for Exon and Exon+Intron counting. We reason that especially very low input material such as from single nuclei sequencing might profit from including reads that potentially originate from nascent RNAs. Another unique feature of *zUMIs* is that it allows for downsampling of reads before collapsing UMIs, thus enabling the user to assess whether a library was sequenced to saturation or whether deeper sequencing is necessary to depict the full mRNA complexity. Furthermore, *zUMIs* is flexible with respect to the length and sequences of the BCs and UMIs, supporting protocols that have both sequences in one read [17, 18, 13, 15, 3, 2, 12] as well as protocols that provide UMI and BC in separate reads [19, 20, 21]. This makes *zUMIs* the only tool that is easily compatible with all major UMI-based scRNA-seq protocols.

**Figure 1.**
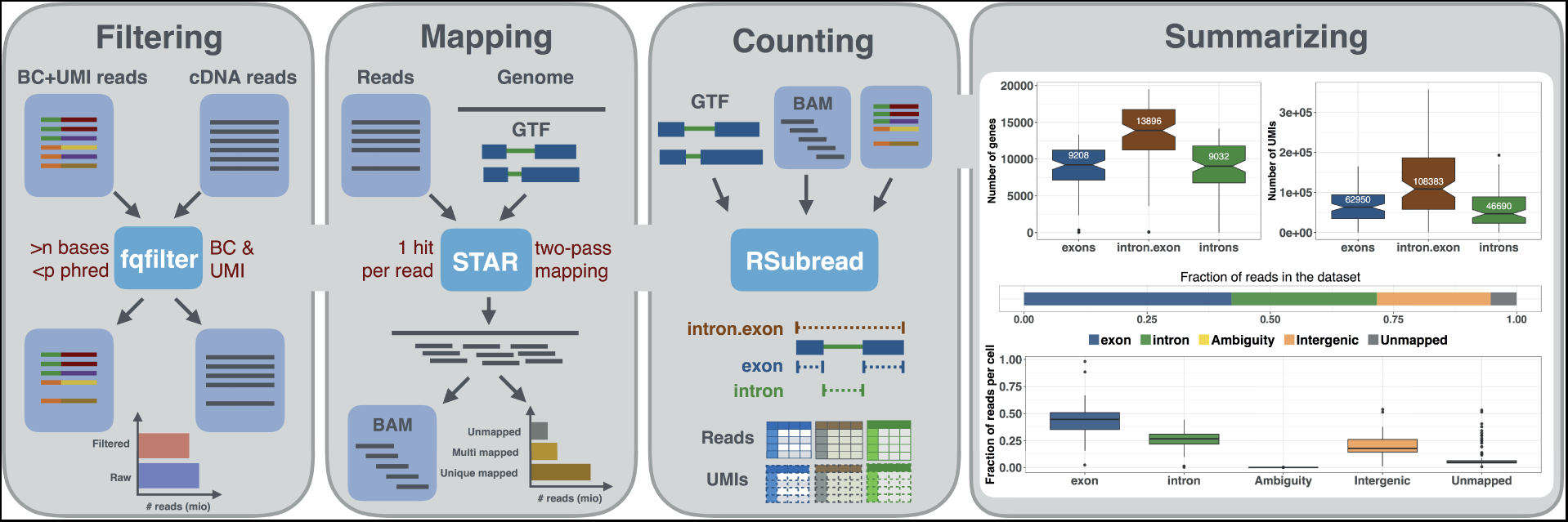
Schematic of the zUMIs pipeline. Each of the grey panels from left to right depicts a step of the *zUMIs* pipeline. First, fastq files are filtered according to user-defined barcode (BC) and unique molecular identifier (UMI) quality thresholds. Next, the remaining cDNA reads are mapped to the reference genome using STAR. Gene-wise read and UMI count tables are generated for Exon, Intron and Exon+Intron overlapping reads. To obtain comparable library sizes, reads can be downsampled to a desired range during the counting step. In addition, *zUMIs* also generates data and plots for several quality measures, such as the number of detected genes/UMIs per barcode and distribution of reads into mapping feature categories.

## Implementation and Operation

### Filtering and Mapping

The first step in our pipeline is to filter reads that have low quality BCs according to a user-defined threshold (Figure 1). This step eliminates the majority of spurious BCs and thus greatly reduces the number of BCs that need to be considered for counting. Similarly, we also filter low quality UMIs.

The remaining reads are then mapped to the genome using the splice-aware aligner STAR [22]. The user is free to customize mapping by using the options of STAR. Furthermore, if the user wishes to use a different mapper, it is also possible to provide *zUMIs* with an aligned bam file instead of the fastq file with the cDNA sequence, with the sole requirement that only one mapping position per read is reported in the bam file.

### Transcript counting

Next, reads are assigned to genes. In order to distinguish Exon and Intron counts, we generate two mutually exclusive annotation files from the provided gtf, one detailing Exon positions, the other Introns. Based on those annotations Rsubread featureCounts [23] is used to first assign reads to Exons and afterwards to check whether the remaining reads fall into Introns, in other words if a read is overlapping with intronic and exonic sequences, it will be assigned to the Exon only. The output is then read into R using data.table [24], generating count tables for UMIs and reads per gene per BC. We then collapse UMIs that were mapped either to the Exon or Intron of the same gene. Note that only the processing of Intron and Exon reads together allows to properly collapse UMIs that can be sampled from the intronic as well as from the exonic part of the same nascent mRNA molecule.

Per default, we only collapse UMIs by sequence identity. If there is a risk that a large proportion of UMIs remains under-collapsed due to sequence errors, *zUMIs* provides the option to collapse UMIs within a given Hamming distance. We compare the two *zUMIs* UMI-collapsing options to the recommended directional adjacency approach implemented in UMI-tools [25], using our in-house example dataset (see Methods). *zUMIs* identity collapsing yields nearly identical UMI counts per cell as UMI-tools, while Hamming distance yields increasingly fewer UMIs/cell with increasing sequencing depth (Figure 2C). Smith et al. [25] suggest that edit distance collapsing without considering the relative frequencies of UMIs might indeed overreach and over-collapse the UMIs. We suspect that this is indeed what happens in our example data, where we find that gene-wise dispersion estimates appear suspiciously truncated as expected if several counts are unduly reduce to one, the minimal number after collapsing (Figure 2D).

**Figure 2.**
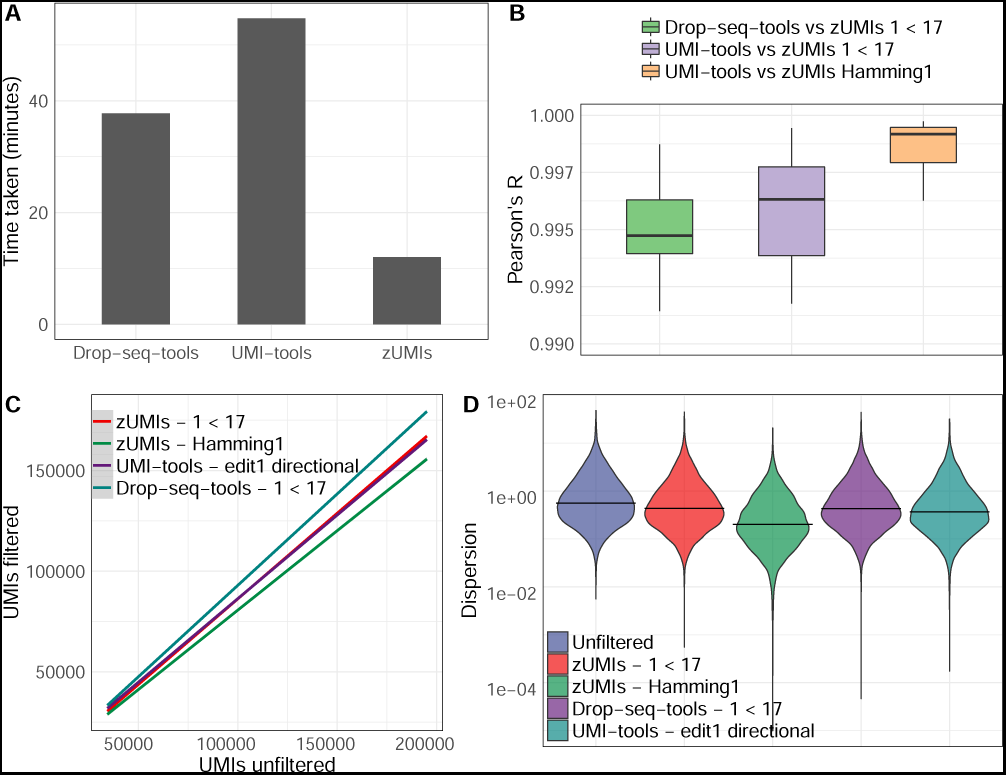
Comparison of different UMI collapsing methods. We compared Drop-seq-tools and UMI-tools with zUMIs using our HEK dataset (227 mio reads). **(A)** Runtime to count exonic UMIs using zUMIs (hamming distance = 0), UMI-tools (“unique” mode) and Drop-seq-tools (edit distance = 0). **(B)** Boxplots of correlation coeficients of gene-wise UMI counts of the same cell generated with different methods. UMI counts generated using zUMIs (quality filter “1 base under phred 17” or hamming distance = 1) were correlated to UMI counts generated using Drop-seq-tools (quality filter “1 base under phred 17”) and UMI-tools (“directional adjacency” mode). **C)** Comparison of the total number of UMIs per cell derived from different counting methods to “unfiltered” counts. **(D)** Violin plots of gene-wise dispersion estimates with different quality filtering and UMI collapsing methods.

However, note that the above described differences are minor. By and large, there is good agreement between UMI counts obtained by UMI-tools [25], the Drop-seq pipeline [13] and *zU-MIs*. The correlation between gene-wise counts of the same cell is > 0.99 for all comparisons (Figure 2B). In light of this, we would consider the > 3 times higher processing speed of *zUMIs* a decisive advantage (Figure 2A).

### Cell Barcode Selection

In order to be compatible with well-based and droplet-based scRNA-seq methods, *zUMIs* needs to be able to deal with known as well as random BCs. As default behavior, *zUMIs* infers which barcodes mark good cells from the data (Figure 3 A,B). To this end, we fit a k-dimensional multivariate normal distribution using the R-package mclust [26, 27] for the number of reads/BC, where k is empirically determined by mclust via the Bayesian Information Criterion (BIC). We reason that only the kth normal distribution with the largest mean contains barcodes that identify reads originating from intact cells. We exclude all barcodes that fall in the lower 1% tail of this kth normal-distribution to exclude spurious barcodes. The HEK dataset used in this paper contains 96 cells with known barcodes and *zUMIs* identifies 99 barcodes as intact, including all the 96 known barcodes. Also for the single-nucleus RNA-seq from Habib et al.[12] *zUMIs* identified a reasonable number of cells: Habib et al. report 10,877 nuclei and zUMIs identified 11,013 intact nuclei. However, we recommend to always check the elbow-plot generated by zUMIs (Figure 3B) to confirm that the cut-off used by zUMIs is valid for a given dataset. In cases where the number of barcodes or barcode sequences are known, it is preferable to use this information. If *zUMIs* is either given the number of expected BCs or is provided with a list of BC sequences, it will use this information and forgo automatic inference.

**Figure 3.**
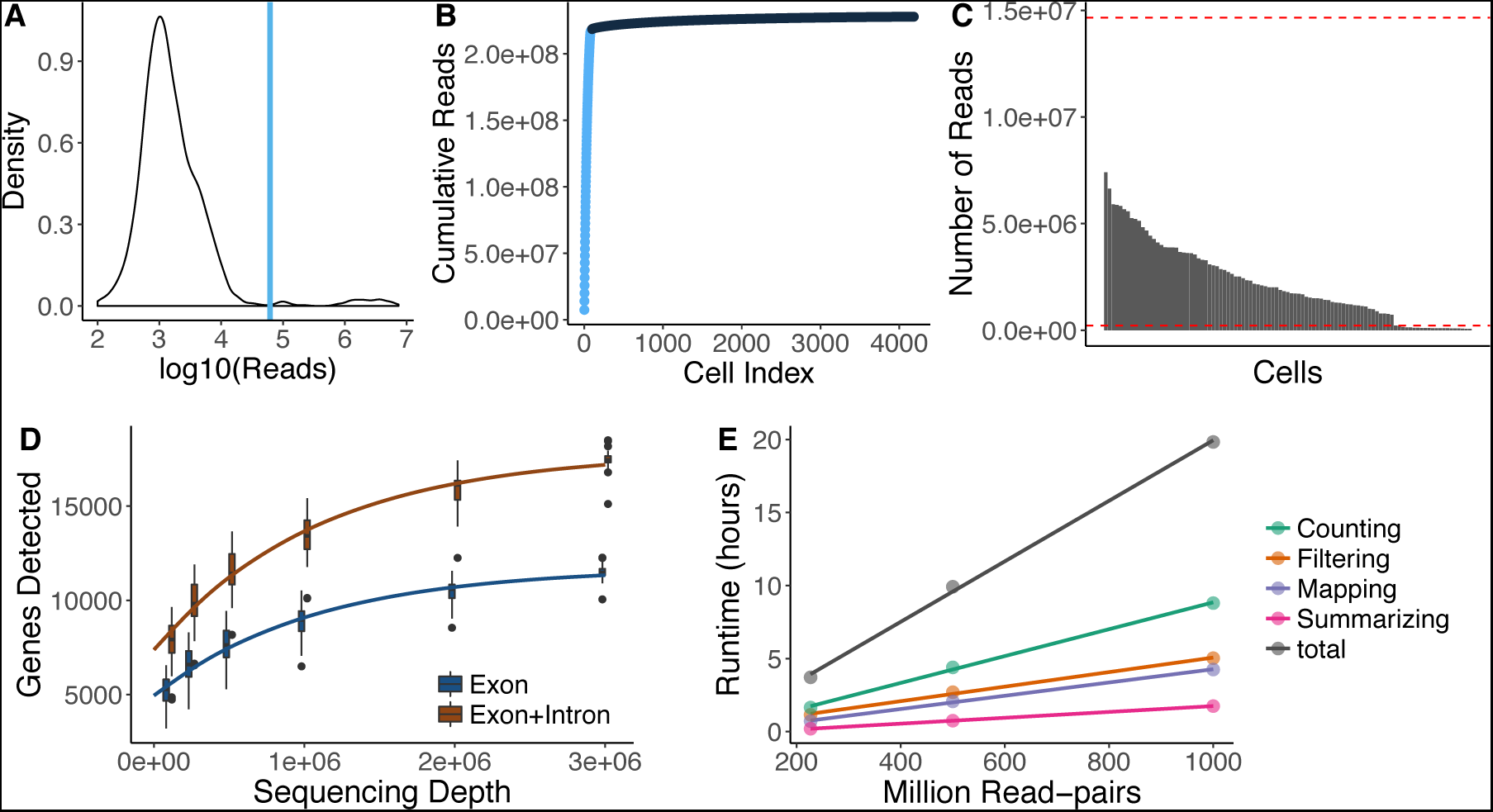
Utilities of zUMIs. Each of the panels shows the utilities of *zUMIs* pipeline. The plots from A-D are the results from the example HEK dataset used in the paper. A) The plot shows a density distribution of reads per barcode. Cell barcodes with reads above the blue line are selected. B) The plot shows the cumulative read distribution in the example HEK dataset where the barcodes in light blue are the selected cells. C) The barplot shows the number of reads per selected cell barcode with the red lines showing upper and lower MAD (Median Absolute Deviations) cutoffs for adaptive downsampling. Here, the cells below the lower MAD have very low coverage and are discarded in downsampled count tables. D) Cells were downsampled to six depths from 100,000 to 3,000,000 reads. For each sequencing depth the genes detected per cell is shown. E) Runtime for three datasets with 227, 500 and 1000 million read-pairs. The runtime is divided in the main steps of the *zUMIs* pipeline: Filtering, Mapping, Counting and Summarizing. Each dataset was processed using 16 threads (“-p 16”).

### Downsampling

scRNA-seq library sizes can vary by orders of magnitude, which complicates normalization [28, 29]. A straight-forward solution for this issue is to downsample over-represented libraries [30]. *zUMIs* has an inbuilt function for downsampling datasets to a user-specified number of reads or a range of reads. By default, *zUMIs* downsamples all selected barcodes to be within three absolute deviations from the median number of reads per barcode (Figure 3 C). Alternatively, the user can provide a target sequencing depth and *zUMIs* will downsample to the specified read number or omit the cell from the downsampled count table if less reads were present. Furthermore, *zUMIs* also allows to specify multiple target read number at once for downsampling. This feature is helpful, if the user wishes to determine whether the RNA-seq library was sequenced to saturation or whether further sequencing would increase the number of detected genes or UMIs enough to justify the extra cost. In our HEK-cell example dataset the number of detected genes starts leveling of at one million reads, sequencing double that amount would only increase the number of detected genes from 9,000 to 10,600, when counting Exon reads (Figure 3D). In line with previous findings [8, 14], the saturation curve of Exon+Intron counting runs parallel to the one for Exon counting, both indicating that a sequencing depth of one million reads per cell is sufficient for these libraries.

### Output and Statistics

*zUMIs* outputs three UMI and three read count tables: gene-wise counts for traditional Exon counting, one for Intron and one for Exon+Intron counts. If a user chooses the downsampling option, 6 additional count tables per target read count are provided. To evaluate library quality, *zUMIs* summarizes the mapping statistics of the reads. While Exon and Intron mapping reads likely represent mRNA quantities, a high fraction of intergenic and unmapped reads indicates low-quality libraries. Another measure of RNA-seq library quality is the complexity of the library, for which the number of detected genes and the number of identified UMIs are good measures (Figure 1). We processed 227 million reads with *zUMIs* and quantified expression levels for Exon and Intron counts on a unix machine using up to 16 threads, which took barely 3 hours. Increasing the number of reads increases the processing time approximately linearly, where filtering, mapping and counting each take up roughly one third of the total time (Figure 3 E). We also observe that the peak RAM usage for processing datasets of 227, 500 and 1000 million pairs was 42 Gb, 89 Gb and 172 Gb, respectively. Finally, *zUMIs* could process the largest scRNA-seq dataset reported to date with around 1.3 million brain cells and 30 billion read pairs generated with 10xGenomics Chromium (see Methods) on a 22-core processor in only 7 days.

### Intron Counting

Recently it has been shown that Intron mapping reads in RNA-seq likely originate from nascent mRNAs and are useful for gene expression estimates [31, 32]. Additionally, novel approaches leverage the ratios of Intron and Exon mapping reads to infer information on transcription dynamics and cell states La Manno et al. [33]. To address this new aspect of analysis, *zUMIs* also counts and collapses Intron-only mapping reads as well as Intron and Exon mapping reads from the same gene with the same UMI. To assess the information gain from intronic reads to estimate gene expression levels, we analyzed a publicly available DroNc-seq dataset from mouse brain ([12], see Methods). For the ∼ 11, 000 single nuclei of this dataset, the fraction of Intron mapping reads of all reads goes up to 61%. Thus, if intronic reads are considered, the mean number of detected genes per cell increases from 1041 for Exon counts to 1995 for Exon+Intron counts. Next, we used the resulting UMI count tables to investigate whether Exon+Intron counting improves the identification of cell types, as suggested in Lake et al. (2016)[11]. The validity and accuracy of counting Introns for single nucleus sequencing methods has recently been demonstrated [34]. Following the Seurat pipeline to cluster cells [35, 36], we find that using Exon+Intron counts discriminates 28 clusters, while we could only discriminate 19 clusters using Exon counts (Figure 4A+B). The larger number of clusters is not simply due to the increase in the counted UMIs and genes. When we permute the Intron counts across cells and add them to the Exon counts, the added noise actually reduces the number of identifiable clusters (Figure 4E).

**Figure 4.**
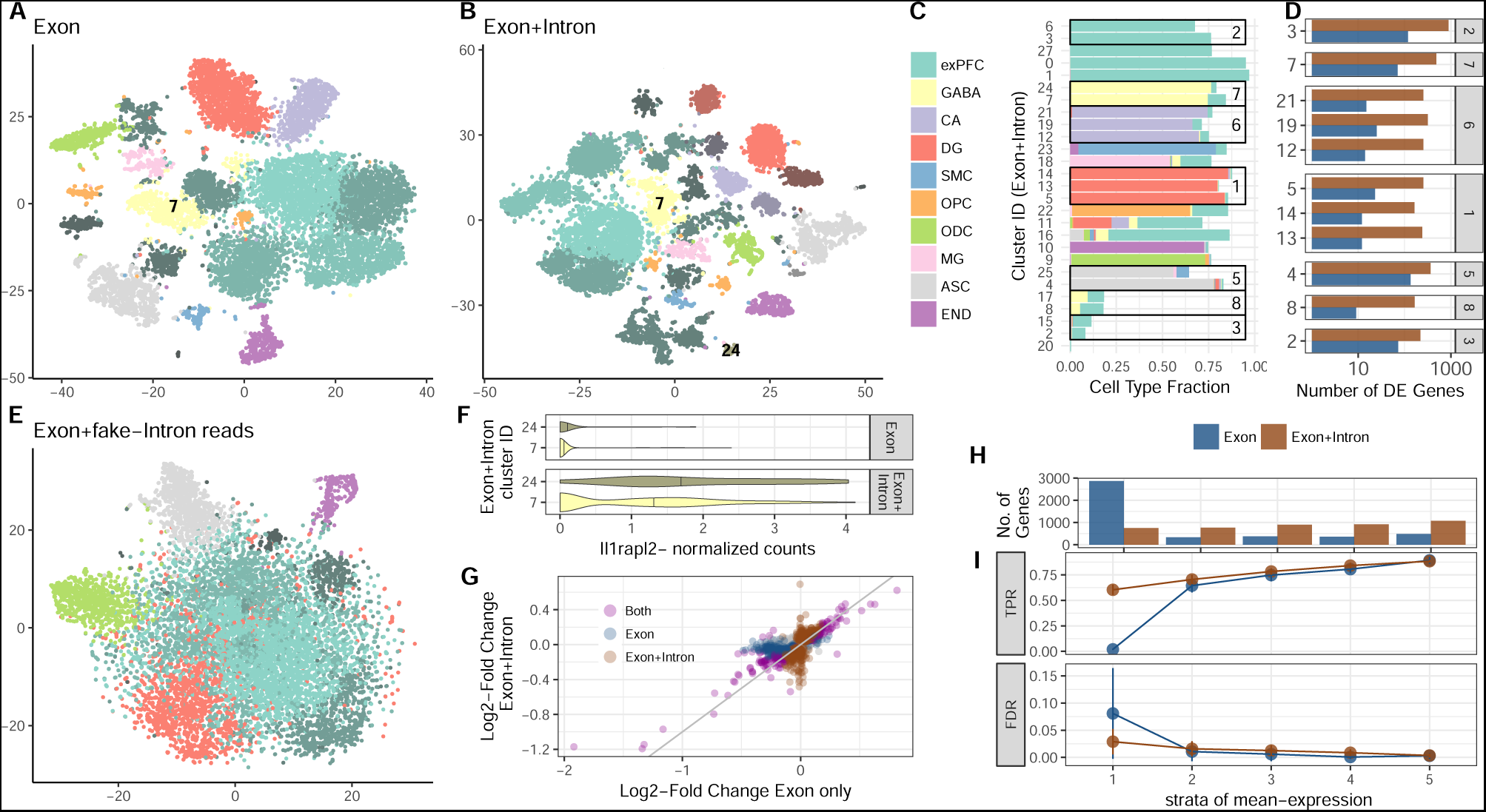
Contribution of Intron reads to biological insights. We analyse published single-nucleus RNA-seq data from mouse prefrontal cortex (PFC) and hippocampus [12] to assess the utility of counting Intron in addition to Exon reads. We processed the raw data with *zUMIs* to obtain expression tables with Exon reads as well as Exon+Intron reads and then use the R-package Seurat [35, 36] to cluster cells. With Exon counts, we thus identify 19 clusters **(A)** and with Exon+Intron counts 27 **(B)**. Clusters are represented as t-SNE plots and colored according to the most frequent cell-type assignment in the original paper [12]: glutamatergic neurons from the prefrontal cortex (exPFC), GABAergic interneurons (GABA), pyramidal neurons from the hippocampal CA region (CA), granule neurons from the hippocampal dentate gyrus region (DG), astrocytes (ASC), microglia (MG), oligodendrocytes (ODC), oligodendrocyte precursor cells (OPC), neuronal stem cells (NSC), smooth muscle cells (SMC) and endothelial cells (END). Different shades of those clusters indicate that multiple clusters had the same major cell-type assigned. If we randomly sample counts from the intron data and add them to the Exon counting, the noise reduces the number of clusters and the Seurat pipeline can only identify 9-11 clusters **(E)**. The composition of each cluster based on Exon+Intron is detailed in panel **(C)** and cells that were not assigned a cell type in Habib et al. [12] are displayed as empty. The boxes mark the clusters that were not split when using Exon data only. For example, cluster 7 from Exon counting that mainly consists of GABAergic neurons, was split into clusters 7, 24 (506, 66 cells) when using Exon+Intron counting. In **(D)**, we show the numbers of genes that were DE (limma p-adj<0.05) between the clusters only found with Exon+Intron counts. The panel numbers represent the Exon counting cluster numbers and the y-axis the Exon+Intron counting cluster number. The log2-fold changes corresponding to these contrasts are also used in **G)**. Among the genes that were additionally detected to be DE by Exon+Intron counting was the marker gene Il1rapl2 (limma p-adj=10^−5^). In **(F)**, we present a violin plot of the normalized counts for Il1rapl2 in cells of the GABAergic subclusters 7 and 24. Log2-fold-changes calculated with Exon+Intron counts correlate well with Exon counts **(G)**. Note that for Exon counting only half as many genes could be evaluated as for Exon+Intron counting and thus only half of the Exon+Intron genes are depicted in **(G)**. Large LFCs are found significant with both counting strategies (purple points are close to the bisecting line). We conduct simulations based on mean and dispersion measured using Exon cluster 0 (1616 cells, ∼ 90% exPFC). In **(I)** we show the expected true positive rate (TPR) and the false discovery rate (FDR) for a scenario comparing 300 vs 300 cells. Results for Exon and Exon+Intron counting were stratified into 5 quantiles according to the mean expression of genes, where stratum 1 contains lowly expressed genes and stratum 5 the most highly expressed genes. The numbers of genes falling into each of the bins using Exon+Intron and Exon counting are depicted in **(H)**.

We continue to further characterize the 7 clusters that were subdivided by the addition of Intron counts (Figure 4D). First, we identify differentially expressed (DE) genes between the newly formed clusters. If we count only Exon reads, there appear to be on average only 10 DE genes between the sub-groups, while Exon+Intron counting yields ∼ 10*×* more DE genes, thus corroborating the signal found with clustering. The log2-fold changes of those additional DE genes estimated with either counting strategy are generally in good agreement, especially large log2-fold changes are detected with both Exon and Exon+Intron counting (Figure 4F). Genes that are detected as DE in only one of our counting strategies have small log2-fold changes and there are more of these small changes detected using Exon+Intron counting.

Detecting more genes naturally increases the chance to also detect more informative genes. Here, we cross-reference the gene list with marker genes for transcriptomic subtypes detected for major cell types of the mouse brain [37] and find that ∼ 5% of the additional genes are also marker genes. Having a closer look at cluster 7, it was split into a bigger (7) and a smaller cluster (24) using Exon+Intron counting (Figure 4A-C), we find one marker gene (Il1rapl2) to be DE between the sub-clusters using Exon+Intron counting, while Il1rapl2 had only spurious counts using Exon counts. Il1rapl2 is a marker for transcriptomic subtypes of GABAergic Pvalb-type neurons [37], suggesting that the split of cluster 7 might be biological meaningful (Figure 4E).

In order to evaluate the power gained by Exon+Intron counting in a more systematic way, we perform power simulations using empirical mean and dispersion distributions from the largest and most uniform cluster (∼ 1500 cells) [9]. For a fair comparison, we include all detected genes and thus there are on average 4*×* more genes in the lowest expression quantile for Exon counting than for Exon+Intron counting (Figure 4H). For those genes, expression is too spurious to be used for differential expression analysis, while for Exon+Intron counting we have on average 60% power to detect a DE gene in the first mean expression bin with a well controlled FDR (Figure 4G). In summary, the increased power for Exon+Intron counting and probably also the larger number of clusters is due to a better detection of lowly expressed genes. Furthermore, we think that, although potentially noisy, the large number of additionally detected genes makes Exon+Intron counting worthwhile, especially for single-nuclei sequencing techniques that are enriched for nuclear nascent RNA transcripts, such as DroNc-seq [12]. Additionally, Exon+Intron counting may help extracting as much information as possible from low coverage data as generated in the context of high-throughput cell atlas efforts (eg 10,000-20,000 reads/cell [38, 39]. Lastly, users should always exclude the possibility of intronic reads stemming from genomic DNA contamination in the library preparation by confirming low intergenic mapping fractions using the statistics output provided by *zUMIs*.

### Conclusion

*zUMIs* is a fast and flexible pipeline processing raw reads to obtain count tables for RNA-seq data using UMIs. To our knowledge it is the only open source pipeline that has a barcode and UMI quality filter, allows Intron counting and has an integrated downsampling functionality. These features ensure that *zUMIs* is applicable to most experimental designs of RNA-seq data, including single nucleus sequencing techniques, droplet-based methods where the BC is unknown, as well as plate-based UMI-methods with known BCs. Finally, *zUMIs* is computationally efficient, user-friendly and easy to install.

## Methods

### Analysed RNA-seq datasets

HEK293T cells were cultured in DMEM High Glucose with L-Glutamine (Biowest) supplemented with 10 % Fetal Bovine Serum (Thermo Fisher) and 1 % Penicillin/Streptomycin (Sigma-Aldrich) in a 37 °C incubator with 5 % CO2. Cells were passaged and split every 2 or 3 days. For single-cell RNA-seq, HEK293T cells were dissociated by incubation with 0.25 % Trypsin (Sigma-Aldrich) for 5 minutes at 37 °C. The single-cell suspension was washed twice with PBS and dead cells stained with Zombie Yellow (Biolegend) according to the manufacturer’s protocol. Single-cells were sorted into DNA LoBind 96-well PCR plates (Eppendorf) containing lysis buffer with a Sony SH-800 cell sorter in 3-drop purity mode using a 100 µm nozzle. Next, single-cell RNA-seq libraries were constructed from one 96-well plate using a slightly modified version of the mcSCRB-seq protocol. Reverse transcription was performed as described previously [40], with the only change being the use of KAPA HiFi HotStart enzyme for PCR amplification of cDNA. Resulting libraries were sequenced using an Illumina HiSeq1500 with 16 cycles in Read 1 to decode cell barcodes (6 bases) and UMIs (10 bases) and 50 cycles in Read 2 to sequence into the cDNA fragment, obtaining ∼ 227 million reads. Raw fastq files were processed using *zUMIs*, mapping to the human genome (hg38) and Ensembl gene models (GRCh38.84).

Furthermore, we anlysed data from 1.3 million mouse brain cells generated on the 10xGenomics Chromium platform [2]. Sequences were downloaded from the NCBI Sequence Read Archive under accession number SRP096558. The data consist of 30 billion read pairs from 133 individual samples. In these data, read 1 contains 16 bp for the cell barcode and 10 bp for the UMI and read 2 contains 114 bp of cDNA. *zUMIs* was run using default settings and we allowed 7 threads per job for a total of up to 42 threads on an Intel Xeon E5-2699 22-core processor. Finally, we obtained mouse brain DroNc-seq read data [12] from the Broad Institute Single Cell Portal (https://portals.broadinstitute.org/single_cell/study/dronc-seq-single-nucleus-rna-seq-on-mouse-archived-brain). This dataset consists of ∼1615 million read pairs from ∼ 11,000 single nuclei. Read 1 contains a 12bp cell barcode and a 8bp UMI and read 2 60bp of cDNA.

The two mouse datasets were mapped to genome version mm10 and applying Ensembl gene models (GRCm38.75).

### Power simulations and DE analysis

We evaluated the power to detect differential expression with the help of the powsimR package [9]. For the DroNc-seq dataset, we estimated the parameters of the negative binomial distribution from one of the identified clusters, namely cluster 0, compromising 1500 glutamatergic neuronal cells from the prefrontal cortex (Figure 4D). Since we detect more genes with Exon+Intron counting (4433 compared to 1782), we included this phenomenon also in our read count simulation by drawing mean expression values for a total of 4433 genes. This means that the table includes sparse counts for the Exon counting. Log2 fold changes were drawn from a gamma distribution with shape equal to 1 and scale equal to 2. In each of the 25 simulation iterations, we draw an equal sample size of 300 cells per group and test for differential expression using limma-trend [41] on log2 CPM values with scran [42] library size correction. The TPR and FDR are stratified over the empirical mean expression quantile bins.

For the differential expression analysis between clusters, we use the same DE estimation procedure as in the simulations: scran normaliztion followed by limma-trend DE-analysis (c.f. [43]).

### Cluster Identiication

After processing the DroNc-seq data [12] with zUMIs as described above, we cluster cells based on UMI counts derived from Exons only and Exons+Introns reads using the Seurat pipeline [35, 36]. First, cells with fewer than 200 detected genes were filtered out. The filtered data were normalized using the ‘LogNormalize’ function. We then scale the data by regressing out the effects of the number of transcripts and genes detected per cell using the ‘ScaleData’ function. The normalized and scaled data are then used to identify the most variable genes by fitting a relationship between mean expression (Exp-Mean) and dispersion (LogVMR) using the ‘FindVariableGenes’ function. The identified variable genes are used for Principle Component Analysis (PCA) and the top 20 PCs are then used to find clusters using graph based clustering as implemented in ‘FindClusters’. To illustrate that the additional clusters found by counting Exon+Intron reads are not spurious, we use Intron-only UMI-counts from the same data to add to the observed Exon only counts. More specifically, to each gene we add scran-sizeFactor corrected Intron counts from the same gene after permuting them across cells. We assessed the cluster numbers from 100 such permutations.

### Comparison of UMI collapsing strategies

In order to validate *zUMIs* and compare different UMI collapsing methods, we used the HEK dataset described above. We ran *zUMIs* (1) without quality filtering, (2) filtering for 1 base under Phred 17 and (3) collapsing similar UMI sequences within a hamming distance of 1. To compare with other available tools, we ran the same dataset using the Drop-seq-tools version 1.13 [13] and quality filter “1 base under Phred 17” without edit distance collapsing. Lastly, the HEK dataset was used with UMI-tools [25] in (1) “unique” and (2) “directional adjacency” mode with edit distance set to 1. Furthermore, we compared the output of *zUMIs* from the DroNc-seq dataset when using default parameters (“1 base under Phred 20”) to UMI-tools in (1). “unique”, (2) “directional adjacency” and (3) “cluster” settings. For each setting and tool combination, we compared per-cell/per-nuclei UMI contents in a linear model fit.

### Availability of Source Code and Requirements

- Project name: zUMIs
- Project home page: https://github.com/sdparekh/zUMIs
- Operating system(s): UNIX
- Programming language: shell, R, perl
- Other requirements: STAR >= 2.5.3a, R >= 3.4, Rsubread >= 1.26.1, pigz >= 2.3 & samtools >= 1.1
- License: GNU GPLv3.0
- Research Resource Identification Initiative ID: SCR_016139

### Availability of supporting data and materials

All data that were generated for this project were submitted to GEO under accession GSE99822.

## Declarations

### List of Abbreviations

scRNA-seq: single-cell RNA-sequencing
UMI: Unique Molecular Identifier
BC: Barcode
MAD: Median Absolute Deviation

## Author’s Contributions

SP and CZ designed and implemented the pipeline. BV tested the pipeline and helped in power simulations. All authors contributed to writing the manuscript.

